# Cysteine restriction induces ferroptosis depending on the polyamine biosynthetic pathway in hepatic cancer cells

**DOI:** 10.1101/2024.02.29.582667

**Authors:** Keisuke Tada, Hironari Nishizawa, Hiroki Shima, Akihiko Muto, Motoshi Wada, Kazuhiko Igarashi

**Affiliations:** Department of Biochemistry, Tohoku University Graduate School of Medicine, Sendai 980-8575, Japan; Department of Pediatric Surgery, Tohoku University Graduate School of Medicine, Sendai 980-8575, Japan

**Keywords:** Cell death, cysteine, ferroptosis, methionine, methionine adenosyltransferase, polyamine, S-adenosylmethionine, sulfur-containing amino acid, liver, hepatoma

## Abstract

**Background and Aims:** Metabolic activities are also known to affect responses and disease processes of the liver which is a central organ for organismal metabolism. Liver diseases such as intestinal failure associated liver disease (IFALD) and hepatocellular carcinoma are known to be affected by nutrition contents, but the mechanisms behind them remain unclear. In this study, we aimed to reveal the relationship between the concentration of sulfur-containing amino acids and hepatocellular response, and further investigated the mechanism focusing on methionine adenosyltransferase (MAT), which plays the central role in methionine metabolism by synthesizing *S*-adenosylmethionine (SAM).

**Methods:** Mouse hepatoma Hepa1 cells were cultured in media with reduced amounts of cysteine, methionine, or both. Cell death was monitored using propidium iodide (PI) and annexin V staining followed by flow cytometry. Inhibitors of ferroptosis (Fer-1), autophagy (GSK872), SAM synthesis (cycloleucine), or polyamine synthesis (sardomozide and DFMO) were used.

**Results:** Cysteine restriction induced marked cell death, whereas simultaneous restriction of cysteine and methionine fully suppressed the cell death. Cysteine restriction-induced cell death was suppressed with Fer-1 and GSK872, suggesting the involvement of ferroptosis in this process. Cysteine restriction-induced cell death was also suppressed by knockdown of MAT2A or its inhibitor cycloleucine. Furthermore, inhibitors of several enzymes in the polyamine biosynthetic pathway also suppressed the cell death. In contrast, primary culture of mouse hepatocytes did not show cell death upon cysteine restriction.

**Conclusions:** These results suggest that SAM-polyamine metabolism is a critical modulator of ferroptosis of hepatic cancer cells. Since normal liver cells were more resistant to ferroptosis than cancer cells, cysteine restriction may be exploited in treating hepatic cancer by inducing ferroptosis specifically in cancer cells without affecting normal cells in the liver.

**Graphical abstrct:** 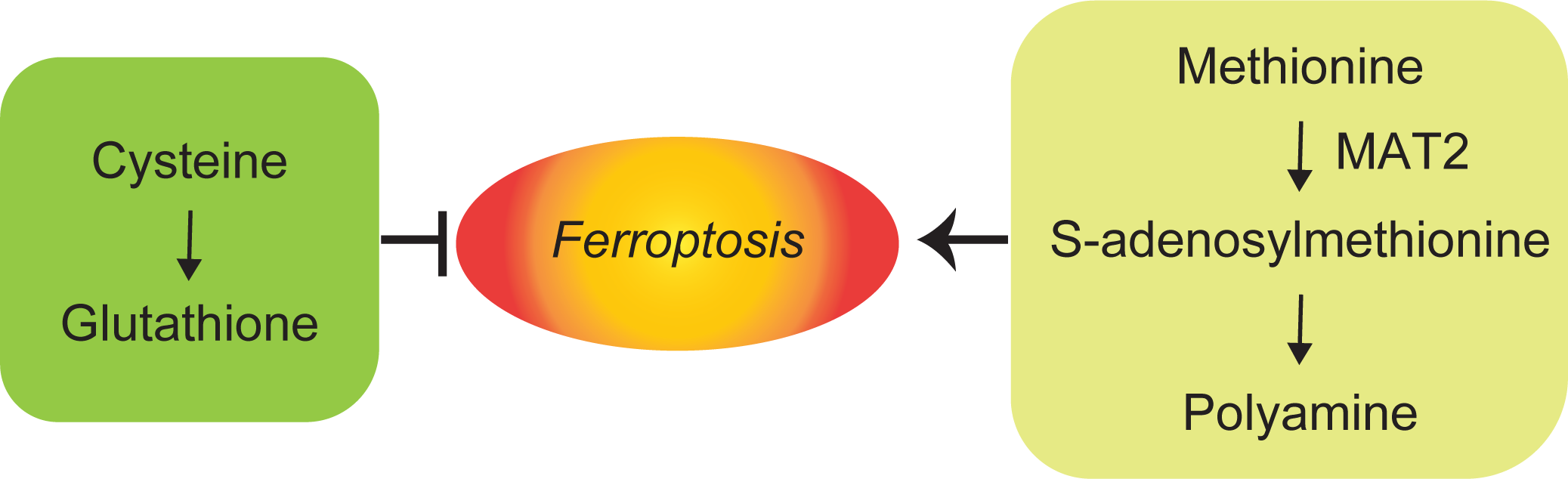

## INTRODUCTION

While the liver is a central organ for organismal metabolism, metabolic activities are also known to affect in turn responses and disease processes of the liver. One of such diseases is intestinal failure-associated liver disease (IFALD) in the field of pediatric surgery (1). IFLAD is a liver disease found in patients who require long-term parenteral nutrition against the background of severe intestinal dysfunction that prevents them from absorbing enough water and nutrients necessary for survival and growth. Because IFALD often progresses to liver cirrhosis (2), control of its onset and progression is very important. As one of the etiologies of IFALD, parenteral nutrition components such as carbohydrates, lipids and amino acids and their administration methods have been pointed out. In addition, various factors have been pointed out, including bacterial infections via parenteral nutrition catheters and translocation of bacteria due to the deterioration of the intestinal epithelial barrier function (3). Although improvements in nutrient administration and improved infection control have resulted in better control of IFALD, it is still not curable. While it has been previously reported that excess methionine causes cholestasis (4), there are few reports suggesting a relationship between excess amino acids and liver damage, and the detailed mechanism remains unclear.

On the other hand, reducing nutritional factors has been shown to affect disease processes of the liver as well (5). The application of methionine restriction to cancer therapy has been discussed in a wide range of cancer types including the liver cancer (6–9). The importance of HNF4α in the therapeutic effect of methionine restriction on liver cancer has been pointed out based on the finding that HNF4α knockdown allowed liver cancer cell lines to tolerate methionine restriction therapy (10). It has been reported that methionine restriction shows a lifespan extension effect in several model systems. Methionine restriction improves glucose metabolism and lipid metabolism in hepatocytes and whole body cells (11). Promoting the catabolism of S-adenosylmethionine (SAM) in Drosophila contributes to lifespan extension due to dietary restriction (12). Regarding the relationship between hepatocytes and amino acids, there are reports that the concentration of extracellular amino acids determines the degree of maturity of hepatocytes (13) and that lack of extracellular amino acids induces hepatocytes into a quiescent state (14).

Methionine adenosyltransferase (MAT) is a central enzyme in methionine metabolism, synthesizing SAM from methionine and ATP (15). The subunits of MAT are encoded by three genes, *Mat1a*, *Mat2a* and *Mat2b*. MAT has multiple isozymes depending on its subunit composition. MAT1 is a tetramer of MAT1A whereas MAT3 is a dimer of MAT1A (16, 17). MAT2 is composed of the catalytic subunit MAT2A and the regulatory subunit MAT2B that stabilizes MAT2A (16–18). MAT1 and MAT3 are expressed in the liver whereas MAT2 is expressed in almost all tissues of the body, developing liver and hepatocellular carcinoma but not in the normal liver (15, 19). Upon transformation of hepatocytes to cancer, the expression of MAT1A is decreased whereas that of MAT2A is increased (20). SAM is a major intracellular methyl group donor, and is used for methylation of DNA, RNA, and histones via methyltransferases, resulting in changes in gene expression (21). Consistent with its roles in gene regulation, both MAT2 and MAT1 are present in the nuclear compartment (22–24). In addition to histones, non-histone proteins such as transcription factors, ribosomal proteins and translation factors are also methylated to regulate their activity (25, 26). Furthermore, SAM is involved in the methylation of lipids and carbohydrates (27, 28). Among various organs and cells, it is known that transmethylation reactions are most actively carried out in the liver (29). SAM is also used in the polyamine synthesis (30). Sulfur-containing amino acids methionine and cysteine are also involved in the synthesis of glutathione (GSH), one of the critical reducing agents against oxidative stress (30).

Considering the possibility that alterations in the sulfur-containing amino acids alter the fate of the liver cells, we investigated the effect of restriction of these amino acids using Hepa1 cells derived from mouse hepatoma and primary mouse hepatocyte culture as cell models. We found efficient induction of ferroptosis upon cysteine restriction in Hepa1 cells but not in primary liver cells. Cysteine restriction-induced ferroptosis of Hepa1 cells was dependent on MAT2 and polyamine synthesis. Our findings suggest a connection of ferroptosis and SAM metabolism in proliferating liver cells.

## RESULTS

### Cysteine restriction-induced cell death is rescued by methionine restriction

To investigate whether the concentrations of methionine and cysteine affect cell death, Hepa-1 cell lines were cultured in culture media containing these amino acids at 10 μM or 200 μM, and the cell death was analyzed by flow cytometry (Table. 1, Fig. 1*A*). Cysteine restriction (i.e., 10 μM) induced significant cell death, whereas methionine restriction did not. Interestingly, simultaneous restriction of cysteine and methionine did not induce cell death (Fig. 1*B-D*). It should be noted that methionine restriction did not reduce cell proliferation (Fig.1*E*). Previous reports have shown that cell death induced by cysteine-free medium is ferroptosis (32–34). In order to determine the type of cell death induced by cysteine restriction in this study, we compared effects of various cell death inhibitors in combination with cysteine restriction (Fig. 2*A*). Fer-1, a ferroptosis inhibitor, markedly inhibited cysteine-restricted cell death, but Z-VAD-fmk, an apoptosis inhibitor, did not significantly suppress cell death. On the other hand, GSK872, an autophagy inhibitor, showed a certain cell death suppression effect, but the effect was smaller than that of Fer-1 (Fig. 2*A*). These results suggest that cysteine restriction-induced cell death of Hepa1 cells is mainly due to ferroptosis, although other type of cell deaths could coexist. The execution of ferroptosis requires the presence of reactive oxygen species (ROS) that leads to lipid peroxidation (35, 36). To examine the effect of ROS on cysteine-restricted cell death, we administered N-acetylcysteine (NAC) which is a ROS scavenger, and quantified cell death (Fig. 2*B*). NAC markedly suppressed cysteine-restricted cell death, suggesting that ROS is involved in cysteine-restrict ion-induced cell death.

**Figure 1.**
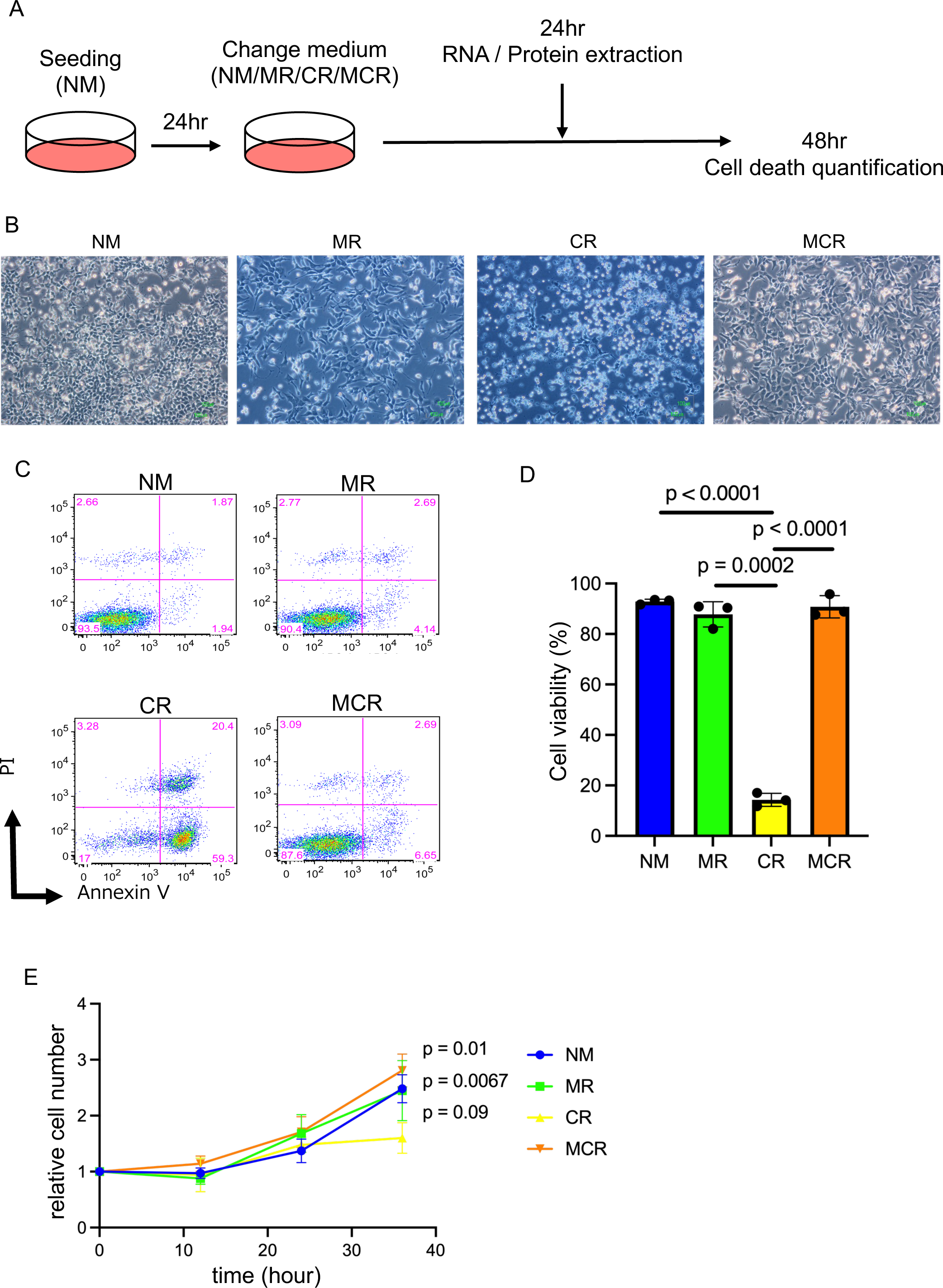
Cysteine restriction-induced cell death is rescued by simultaneous methionine restriction. (A) Experimental flow chart. Normal medium (NM), methionine restriction (MR), cysteine restriction (CR), and methionine and cysteine restriction (MCR). (B, C, D) Light microscope image (B) and percentage of live cells (C, D) of Hepa-1 cells cultured under the indicated conditions for 48 hours. Representative data (B, C) or means and SD (D) of three experiments are shown. (E) Proliferation Hepa-1 cells under the indicated conditions. Means and SD of three experiments are shown.

**Figure 2.**
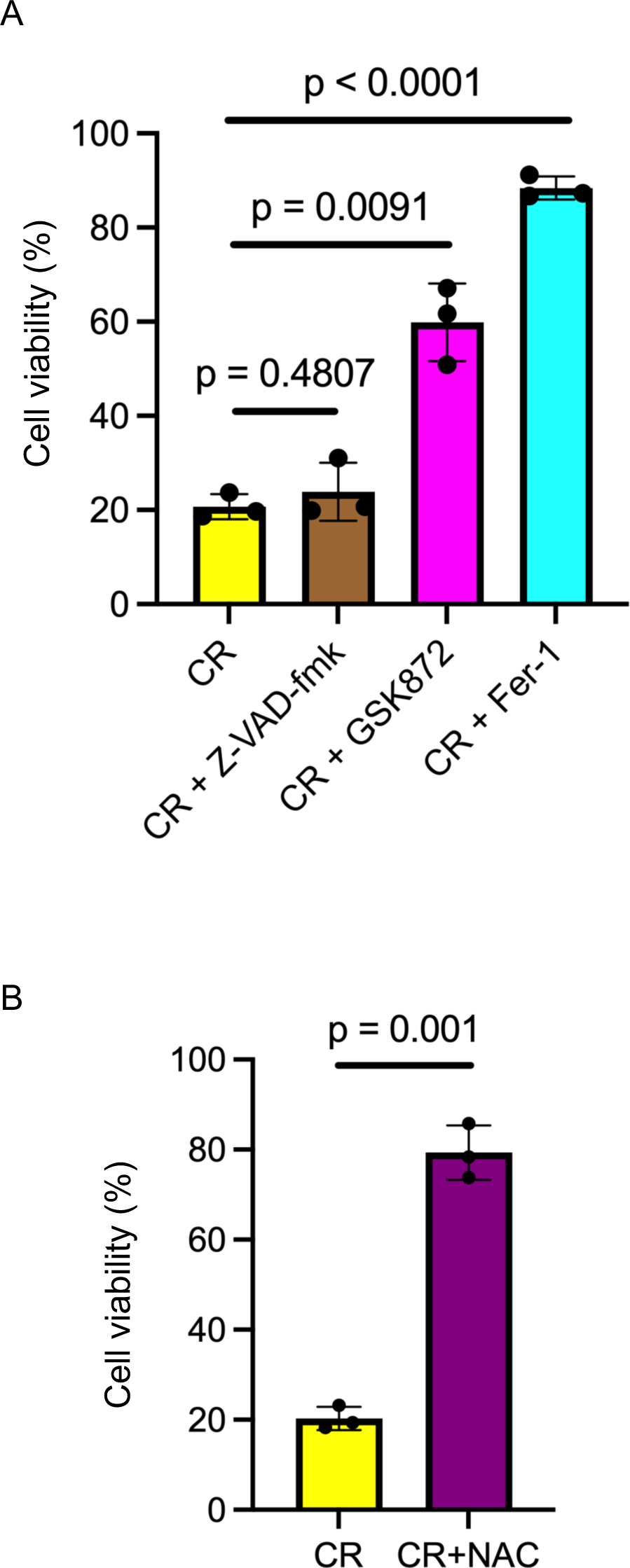
cysteine restriction-induced cell death is ferroptosis. (A) Percentage of live cells in Hepa-1 cells cultured for 48 hours in cysteine-limited medium supplemented in the presence of Z-VAD-fmk, GSK872 or Fer-1. (B) Percentage of liive cells in Hepa-1 cells cultured for 48 hours in cysteine-limited medium supplemented with NAC. Means and SD of three experiments are shown.

### Cysteine restriction-induced cell death depends on SAM synthesis

In addition to contributing to protein synthesis, methionine is an important SAM precursor. It has been reported that homeostasis of SAM synthesis is maintained by feedback control by SAM (37, 38). Therefore, we next evaluated the gene expression of *Mat2a* and *Mat2b* involved in SAM synthesis and the amounts of the MAT2A and MAT2B proteins (Fig. 3). *Mat2a* mRNA was upregulated under methionine restriction as expected, but not under cysteine restriction (Fig. 3*A*). In addition, an increase in the *Mat2a* mRNA by methionine restriction was also observed under cysteine restriction. On the other hand, *Mat2b* mRNA expression was elevated by both methionine and cysteine restrictions. The MAT2A protein under methionine restriction showed changes that mirrored its mRNA, but a milder increase was also observed under cysteine restriction (Fig. 3*B* and *C*). There was no significant difference in the amount of the MAT2B protein under these conditions. These results suggested that increased MAT2A may protect against cell death induced by cysteine restriction.

**Fig 3.**
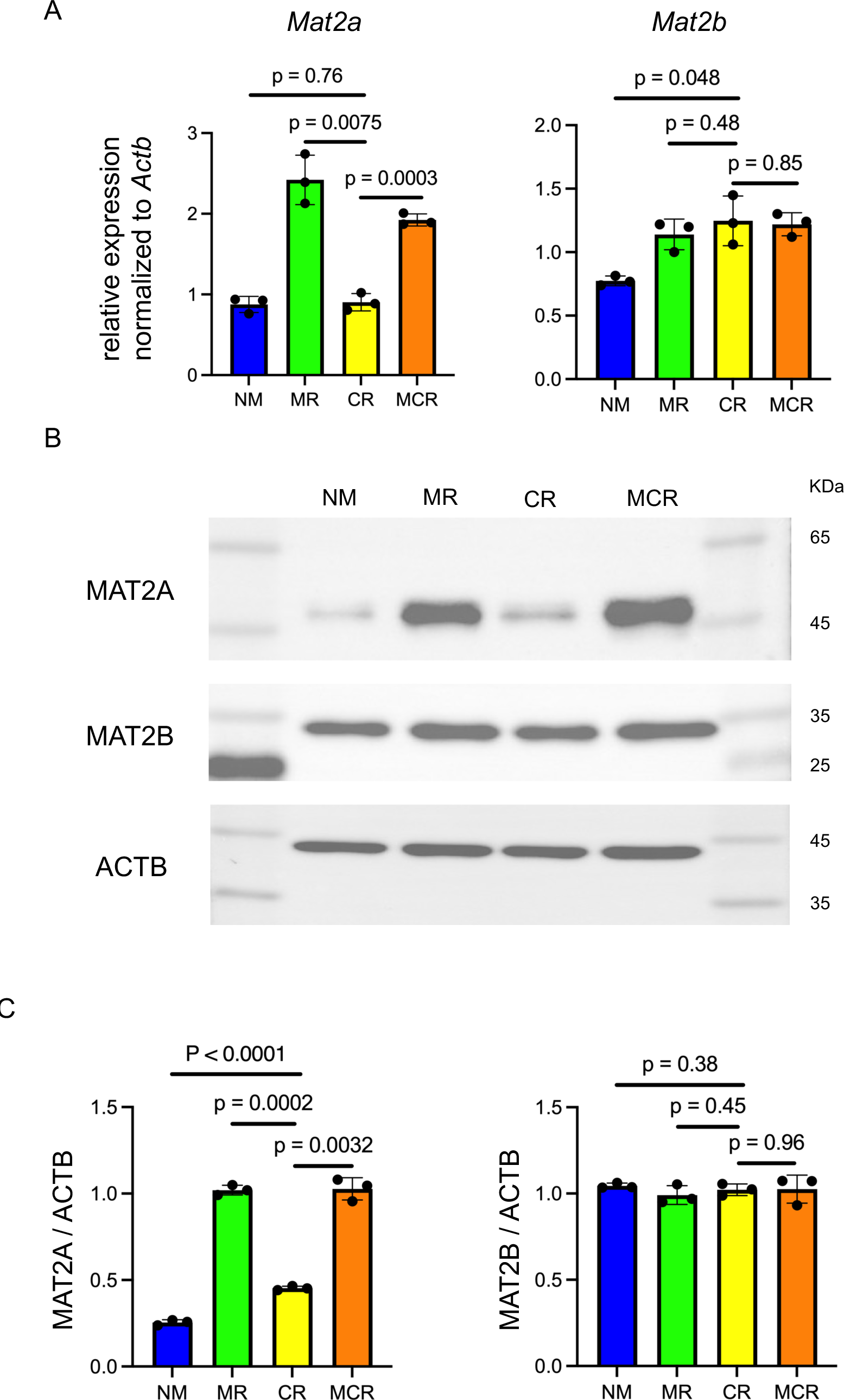
Changes of MAT2A expression under restriction of sulfur-containing amino acids. (A) Expression levels of Mat2a and Mat2b mRNA in Hepa-1 cells cultured for 24 hours under indicated conditions. Relative values were calculated using Actb as an internal control. (B, C) Protein levels of MAT2A, MAT2B, and beta-actin (B) in Hepa-1 cells cultured as above and relative values using beta-actin as an internal control (C). These samples were derived from the same experiment and that gels/blots were processed in parallel. Relative values were calculated by calculating brightness using ImageJ. Means and SD (A, C) and representative data (B) of three experiments are shown.

In order to investigate this possibility, we combined a MAT inhibitor cycloleucine (Cleu) or a siRNA-mediated *Mat2a* knockdown together with cysteine restriction (Fig. 4*A, B*). Surprisingly, cysteine-restricted cell death was not enhanced under these treatments, but cell death was almost completely suppressed. It was suggested that MAT2A acted executively rather than protectively in the process of cell death upon cysteine restriction. Next, to investigate the relationship between changes in cell death caused by cysteine restriction and SAM synthesized by MAT, intracellular SAM levels were measured using an ELISA kit (Fig. 4*C, D*). No significant difference was observed in the intracellular SAM level at 24 hours under any condition, whereas a significant increase in intracellular SAM level at 36 hours was observed under the simultaneous restriction of cysteine and methionine.

**Fig 4.**
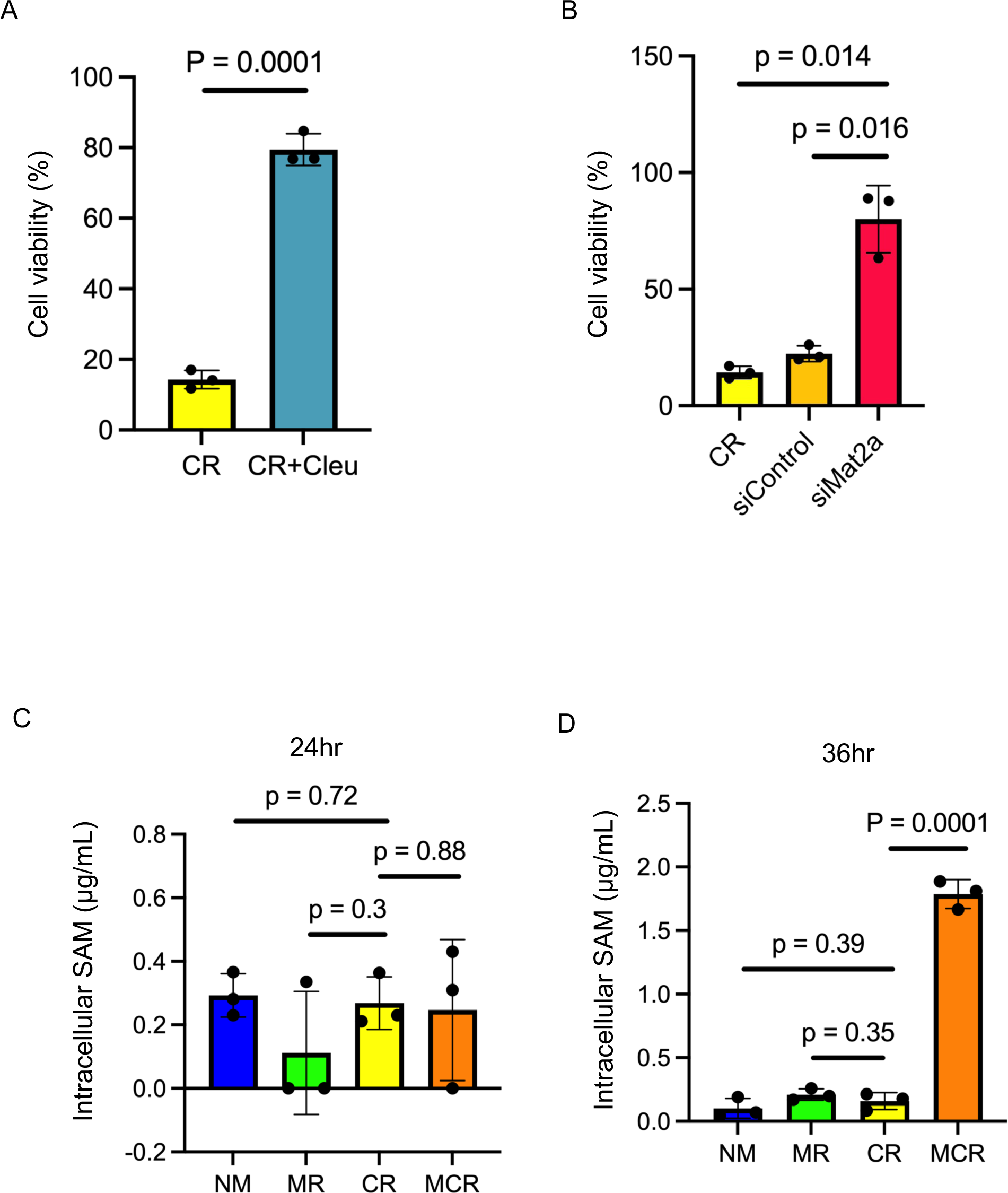
Cysteine restriction-induced cell death depends on SAM synthesis. (A) Percentage of live cells in Hepa-1 cells cultured for 48 hours in cysteine-restricted medium supplemented with or without cycloleucine. (B) Percentage of live cells in *Mat2a*-knockdown Hepa-1 cells cultured as above. (C, D) Endogenous SAM measurements in Hepa-1 cells cultured for 24 hours (C) or 36 hours (D) under indicated conditions. Means and SD of three experiments are shown.

### Cysteine restriction-induced cell death is rescued by inhibition of polyamine biosynthesis

Next, we focused on SAM metabolism to verify the mechanism of cell death due to cysteine restriction (Fig. 5*A*) (39). SAM is converted to SAH by methyltransferases such as GNMT, and then metabolized to homocysteine. Homocysteine is synthesized to methionine by methionine synthase, or is transferred to the transsulfuration pathway to be used for the synthesis of cysteine and glutathione (Fig. 5*A*). On the other hand, SAM is also metabolized in the polyamine biosynthetic pathway, converted to decarboxy SAM by SAM decarboxylase (SAMDC), which is then used along with putrescine in the synthesis of spermidine and spermine (Fig. 5*A*). It has been reported that a large amount of ROS is produced during the process of polyamine biosynthesis, and activation of polyamine biosynthesis increases the demand for cysteine in cancer cells (40). To evaluate the effects of the polyamine biosynthetic pathway on cell death during cysteine restriction in this study, we quantified cell death using sardomozide, the SAMDC inhibitor, and DFMO, the ornithine decarboxylase (ODC) inhibitor (Fig. 5*B*). Both inhibitors inhibited cell death under cysteine restriction, suggesting that the polyamine biosynthetic pathway is involved in the execution of this cell death. To investigate the possibility that polyamines themselves induce cell death, putrescine, spermidine, and spermine were administered under the cysteine/methionine simultaneous restriction conditions to evaluate cell death (Fig. 5*C*). Putrescine administration did not cause cell death, whereas spermidine and spermine administration induced significant cell death. However, spermidine- or spermine-induced cell death was not suppressed by Fer-1 (Fig. 5*D*). Unlike endogenous products from the polyamine biosynthetic pathway, exogenous spermine and spermidine appeared to cause cell death other than ferroptosis when added extracellularly. These results suggested that endogenous and exogenous polyamines have distinct roles in cell death.

**Fig 5.**
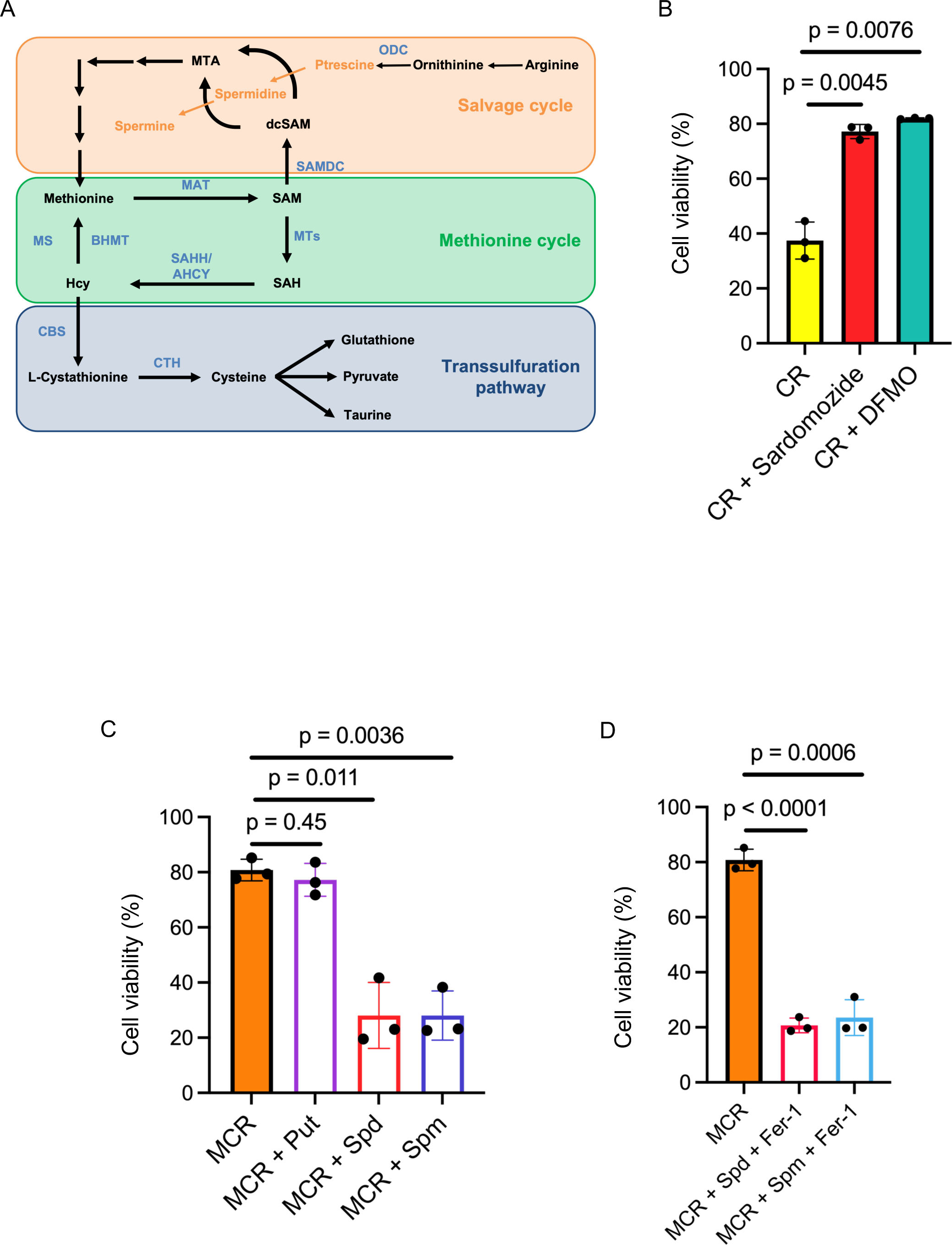
Cysteine restriction-induced cell death is suppressed by inhibition of polyamine biosynthesis. (A) Conceptual diagram of SAM metabolism. BHMT; betaine homocysteine methyltransferase, CBS; cystathionine beta-synthase, CTH; cystathionine gamma-lyase, MAT; methionine adenosyltransferase, MS; methionine synthase, MTA; 5’-methylthioadenosine, MTs; methionine transferase, ODC; ornithine decarboxylase, SAHH/AHCY; S-adenosylhomocysteine hydrolase / adenosylhomocysteinase, SAMDC; S-adenosylmethionine decarboxylase. (B) Percentage of live cells in Hepa-1 cells cultured for 48 hours in cysteine-restricted medium supplemented with Sardomozide or DFMO. (C) Percentage of live cells of Hepa-1 cell line cultured for 48 h in cysteine- and methionine-restricted medium supplemented with putrescine, spermidine, or spermine. (D) Percentage of live cells in Hepa-1 cells cultured as above with Fer-1. Means and SD of three experiments are shown.

### Cysteine restriction does not induce cell death in mouse primary hepatocytes

Next, we investigated how the concentrations of methionine and cysteine affect primary hepatocytes. Primary hepatocytes were extracted from wild type mice by the previously reported method (31). Primary hepatocytes were cultured in the same manner as Hepa1, and cell death was analyzed with flow cytometry (Fig. 6*A, B*). In contrast to Hepa1 cells, the primary hepatocytes did not undergo cell death under cysteine restriction. We then assessed gene expression of *Mat1*, *Mat2a*, *Mat2b* and the MAT1/3, MAT2A, and MAT2B proteins under these conditions (Fig. 7*A-C*). Mat2a mRNA expression tended to increase under methionine restriction, and Mat2b mRNA expression tended to increase under methionine-cysteine simultaneous restriction. Mat1 mRNA expression was very low irrespective of the conditions, suggesting that the effect of pre-culture for 24 hours decreased its expression. There was no significant difference in the protein levels of MAT1/3, MAT2A, and MAT2B under any condition (Fig. 7*B* and *C*), unlike the Hepa1 cells.

**Fig 6.**
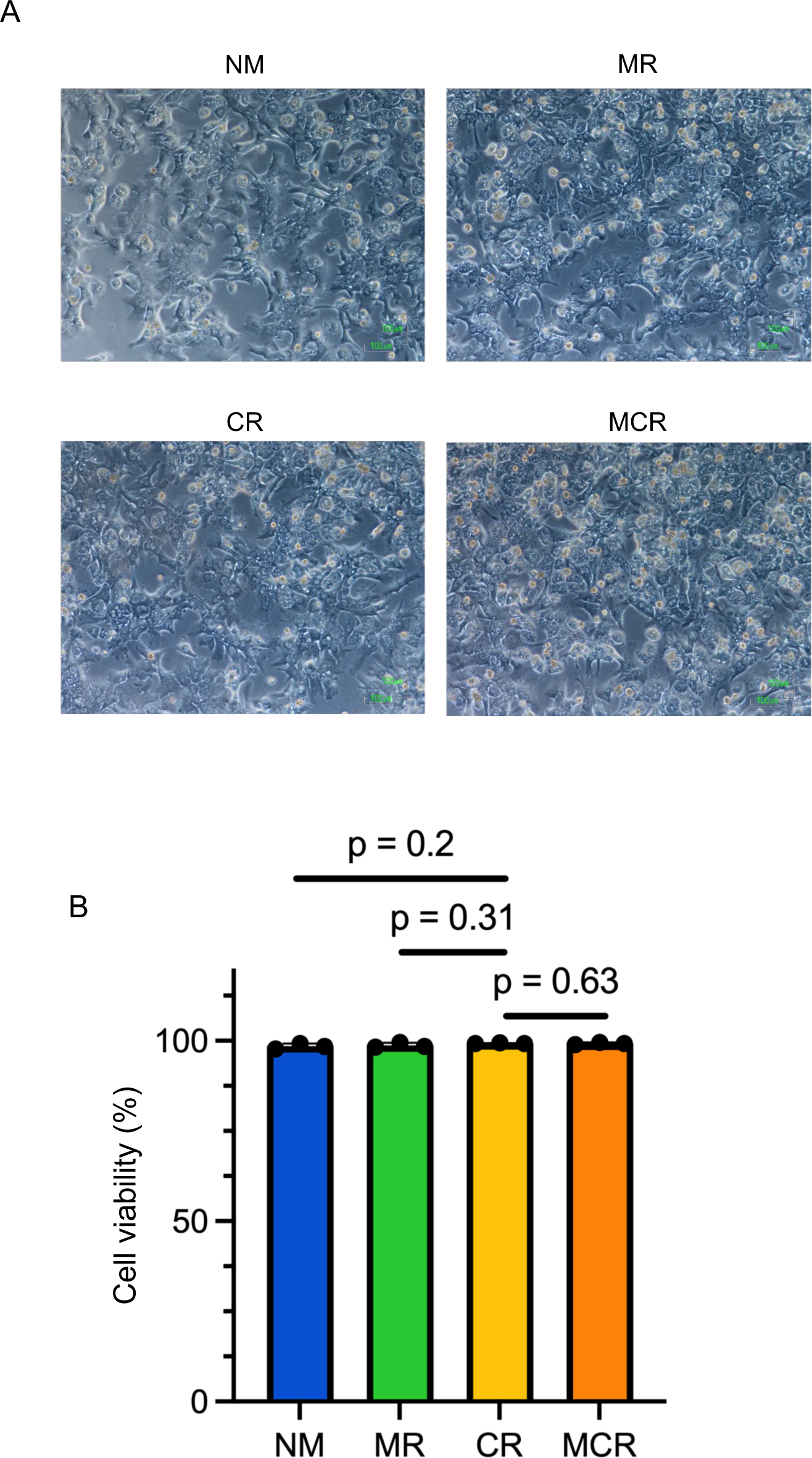
Cysteine restriction does not induce cell death in mouse primary hepatocytes. (A, B) Light microscope image (A) and percentage of live cells (B) of primary hepatocytes cultured for 48 hours under indicated conditions. Representative data (A) or means and SD of three experiments are shown.

**Fig 7.**
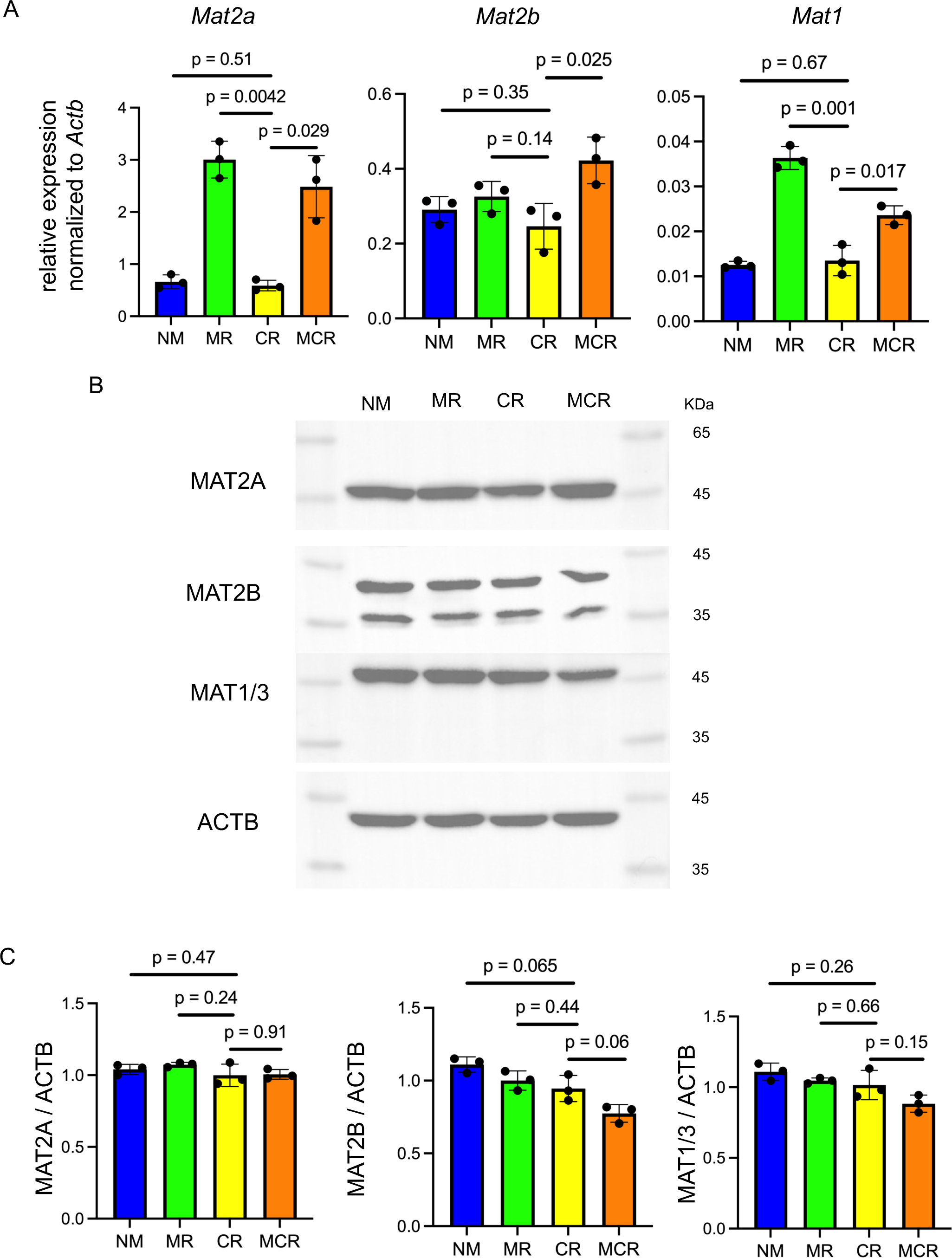
Effects of sulfur-containing amino acid concentrations on MAT expression in primary hepatocytes. (A) Expression levels of Mat1, Mat2a and Mat2b mRNA in mouse primary hepatocytes cultured for 24 hours under indicated conditions. Relative values were calculated using Actb as an internal control. (B, C) Protein levels of MAT1/3, MAT2A and MAT2B, and beta-actin in primary hepatocytes cultured as above (B). Relative values of indicated proteins are shown using beta-actin as an internal control (C). These samples were derived from the same experiment and gels/blots were processed in parallel. Relative values were calculated by calculating brightness using ImageJ. Means and SD (A, C) or representative data (B) of three experiments are shown.

## DISCUSSION

Links of metabolism and liver cell responses are emerging as critical mechanisms in disease conditions including IFALD, cholestasis, life span extension, and cancer therapy (1, 4, 9, 11). Methionine restriction has been reported to extend life span in several model animals (12, 41–45). On the other hand, cystine/cysteine restriction have been shown to promote ferroptosis in several model cell systems (46, 47). However, there have been no reports examining the relationship between cysteine and methionine metabolism in hepatocellular response or its damage. In this study, we found that cysteine restriction efficiently induced cell death in Hepa1 cells derived from murine hepatocarcinoma (Fig. 1*D*). This cell death is judged to be ferroptosis, since it was efficiently suppressed by Fer-1 ferroptosis inhibitor (Fig. 2*A*). GSK872, an autophagy inhibitor, also suppressed the observed cell death (Fig. 2 *A*), suggesting that autophagy is involved in this cell death. A previous report has shown that ferritin-specific autophagy, ferritinophagy, is the cause of cell death in response to cysteine restriction (48). Ferrous iron released upon ferritinophagy has been shown to promote ferroptosis (49). Indeed, ferrous iron is increased during the induction of ferroptosis (50). Consistent with these observations, inhibition of ferritinophagy suppresses ferroptosis (51). Taken together with these previous reports, the present results suggest that Hepa1 cells undergo ferroptosis upon cysteine restriction.

We found that ferroptosis of Hepa1 cells induced by cysteine restriction was completely rescued by simultaneous restriction of methionine. A previous report has shown that methionine is essential for ferroptosis of fibrosarcoma cells induced by cysteine depletion (52). Another report described that Hela cells undergo ferroptosis upon cysteine depletion which is relieved by methionine depletion (32). However, unlike these previous reports, methionine and cysteine were not depleted from the medium in this study and were set at 10 μM. While cell cycle arrest upon methionine depletion was suggested as a mechanism that suppresses cell death upon cysteine depletion (32), this appears not to be the case in our study since Hepa1 cells continued to proliferate even under the methionine-restricted conditions. Our results suggest that the suppression of cell death by simultaneous restriction of cysteine and methionine in this study involves a mechanism other than cell cycle arrest. Surprisingly, both administration of MAT inhibitor and knockdown of *Mat2a* almost completely suppressed the cell death induced by cysteine restriction, suggesting that MAT2 is required for the execution of cell death under cysteine restriction. However, one observation did not agree with this interpretation; when cysteine and methionine were restricted simultaneously, a more pronounced increase in intracellular SAM was observed than when methionine alone was restricted. The cause of this is unknown at present, but one possibility is that SAM consumption which is required for ferroptosis is reduced when cysteine and methionine are simultaneously restricted. Methionine restriction may reduce SAM-dependent reactions leading to ferroptosis. Alternatively, it is possible that lower concentrations of SAM promote ferroptosis by supporting SAM-dependent reactions, whereas higher concentrations of SAM inhibit ferroptosis by supporting the other SAM-dependent reactions. Understanding the mechanism of SAM increase upon simultaneous cysteine and methionine restriction may be important in clarifying these questions.

Our current results suggest that one of the SAM-dependent reactions leading to ferroptosis is polyamine synthetic pathway. There are several reports showing that the activity of the polyamine biosynthetic pathway increases the demand for cysteine in cancer cells (53–56). Importantly, the activity of the polyamine biosynthetic pathway produces large amounts of ROS (40) which can promote ferroptosis (57). We found that inhibition of the two enzymes in polyamine biosynthetic pathway suppressed cysteine restriction-induced cell death. While extracellular administration of polyamines induced cell death, Fer-1 did not suppress this cell death. When taken together, these results suggest that polyamine biosynthetic pathway activity is important for the execution of cysteine restriction-induced cell death but, when added extracellularly, polyamine can induce other types of cell death. It is possible that ROS which is derived from polyamine synthesis promotes ferroptosis whereas excess amounts of polyamines stimulate other types of cell death.

Since polyamines are essential for cell proliferation and are increased in cancer cells (53–55), strategies targeting inhibition of polyamine biosynthesis and depletion of polyamines have been suggested (56). However, our present results raise the possibility that inhibition of polyamine metabolism may actually be beneficial for cancer cells in evading ferroptosis. In contrast, cysteine restriction may be used to induce ferroptosis in the treatment of hepatocellular carcinoma because cysteine restriction did not induce ferroptosis of primary hepatocytes. By adjusting the amounts of cysteine and methionine in cancer microenvironment, it may become possible to induce ferroptosis specifically in cancer cells without killing other types of cells like normal cells and immune cells.

It has been reported that ferroptosis is a trigger for the onset and progression of NAFLD and NASH (58). There are a series of reports linking liver damage and ferroptosis (5, 59–62). In this study, it was suggested that the concentration of sulfur-containing amino acids can alter the metabolism of SAM. Combined with the above reports, we surmise that various liver disorders involve changes in sulfur-containing amino acids and SAM metabolism. Recently, it was pointed out that the expression of ferroptosis-related genes is elevated in a rat IFALD model (63). Taken together with our present observations, it is possible that sulfur-containing amino acids and SAM metabolism are involved in the onset and progression of IFALD.

## Acknowledgements

We thank members of the Departments of Biochemistry, Tohoku University Graduate School of Medicine for discussions and support; the Biomedical Research Core of Tohoku University Graduate School of Medicine for technical support; Institute for Animal Experimentation of Tohoku University Graduate School of Medicine for breeding mice.

## EXPERIMENTAL PROCEDURES

### Animal Experimentation

The wild type mice on the C57BL/6J background were bred at the animal facility of Tohoku University. Mice were housed under specific pathogen-free conditions. All experiments performed in this study were approved by the Institutional Animal Care and Use Committee of the Tohoku University Environmental & Safety Committee.

### Primary hepatocyte isolation

Primary hepatocyte isolation from wild type mice (14 weeks old, male) was performed as previously described (31). Briefly, after anesthesia with isoflurane, the portal vein was cannulated. After cutting the inferior vena cava, the liver was perfused with Liver perfusion medium (Thermo Fisher Scientific) at 37℃ for 10 minutes. Subsequently, after perfusion with HEPES buffer containing 0.05% collagenase for an additional 10 minutes, the liver tissue was excised. The liver was suspended in DMEM and passed through a 70 μm nylon mesh. Centrifugation was performed using Percoll to purify the hepatocytes.

### Cell culture

Mouse hepatoma cell line Hepa1c1c7 (Hepa1) was cultured using DMEM (Dulbecco’s modified Eagle medium) supplemented with 10% FBS, 100 U/ml penicillin, and 0.1 mg/ml streptomycin as a normal medium. Conditioned media for methionine and cysteine were prepared as follows: 10% dialyzed FBS (Gibco, Carlsbad, USA), 100 U/ml penicillin, 0.1 mg/ml streptomycin and 4 mM glutamine (Gibco, Carlsbad, USA) were added in DMEM without glutamine, methionine and cysteine (Gibco, Carlsbad, USA). L-methionine (Sigma-Aldrich, St. Louis, USA) and L-cystine (Sigma-Aldrich, St. Louis, USA) were added to this medium and adjusted to 10 µM or 200 µM, respectively.

### Reagents

Erastin, (1S, 3R)-RSL3 (RSL3), dimethyl sulfoxide (DMSO), α-tocopherol (α-Toc), Ferrostatin-1 (Fer-1), chloroquine, and RNase A were purchased from Sigma-Aldrich (St. Louis, MO, USA). MG132 was purchased from Calbiochem (San Diego, CA, USA). Trypsin was purchased from GL Science (Fukushima, Japan). APC-Annexin V was purchased from Becton, Dickinson and Company (BD) (Franklin Lakes, NJ, USA). Sardomozide dihydrochloride and DFMO were from Axon Medchem (Reston, VA, USA) and were used as inhibitors of polyamine synthesis.

### RNA interference

All siRNAs (siControl: Stealth RNAiTM siRNA Negative Control, Med GC, siMat2a #911: MSS232488) were obtained from Invitrogen (Carlsbad, CA, USA). The target sequence of the siMat2a (911) RNAi is; 5’-AGGAAAGGATTATACCAAAGTGGAC-3’. Hepa1 cells were transfected with siRNAs using Lipofectamine RNAiMAX (Invitrogen) according to the manufacturer’s protocols. After transfection, Hepa1 cells were passaged to dishes or culture plate with culture medium.

### Western Blotting

Cells were treated with trypsin, pelleted, and washed twice in PBS. Cells were lysed by RIPA buffer (50 mM Tris-HCl (pH = 7.4), 1 mM ethylenediaminetetraacetic acid, 150 mM NaCl, 1% (v/v) NP-40, 0.5% (w/v) sodium deoxycholate, 0.1% (w/v) sodium dodecyl sulfate ( SDS)) and then mixed by shaking for 10 min at 4°C. After that, centrifugation was performed at 4°C and 15,000 rpm for 10 minutes, and the supernatant was collected. 5×SDS sample buffer (312.5 mM Tris-HCl (pH = 6.8), 25% (v/v) 2-Mercaptoethanol, 10% (w/v) SDS, 50% (w/v) glycerol, 0.01% (w/v) bromophenol blue; BPB) was added and heated for 5 minutes. Lysates were resolved on 10% SDS–PAGE gels and transferred to PVDF membranes (Millipore, Billerica, MA, USA). The membranes were blocked for 1 hour in blocking buffer [3% Skim milk (Sigma-Aldrich) in T-TBS buffer (0.05% Tween 20 (Sigma-Aldrich) in TBS (tris-buffered saline))] and subsequently incubated with the primary antibodies in T-TBS buffer containing 1% Skim milk overnight at 4°C. After washing in T-TBS, secondary antibodies were incubated with the membranes as above. Immune complexes were detected using Pierce^TM^ ECL Plus Western Blotting Substrate (Thermo Fisher Scientific 32132) and images were captured using ChemiDoc^TM^ MP Imaging System(Bio-Rad). The antibody for ACTB (ab3280) was purchased from Abcam (Cambridge, UK). The antibody for MAT1/3 (sc-28029) was purchased from Santa Cruz Biotech (Dallas, Texas, USA). The antibodies for MAT2A and MAT2B were previously described(23). For the quantification of signals, all samples to be compared were run on the same gel. Bands were quantified using ImageJ. All bands to be compared were quantified on the same image and were within the linear range of detection of the software.

### Quantitative PCR with reverse transcription

Total RNA was purified with RNeasy plus mini kit (Qiagen, Hilden, Germany). Complementary DNA was synthesized by High Capacity cDNA Reverse Transcription Kits (Applied Biosystems, Foster city, CA, USA). Quantitative PCR was performed using LightCycler Fast Start DNA Master SYBR Green I, and LightCycler 96 (Roche). mRNA transcript abundance was normalized to that of *Actb*. Sequences of the qPCR primers are described in Table 2.

**Table 1:** Sulfur-containing amino acid concentration in the culture medium. Sulfur-containing amino acid concentration (μM) used in this study and the name of each conditioned medium are shown.

| Name of medium | Methionine | Cysteine |
| --- | --- | --- |
| Normal Medium; NM | 200 | 200 |
| Methionine restriction; MR | 10 | 200 |
| Cysteine restriction; CR | 200 | 10 |
| Methionine Cysteine restriction; MCR | 10 | 10 |

**Table 2:**
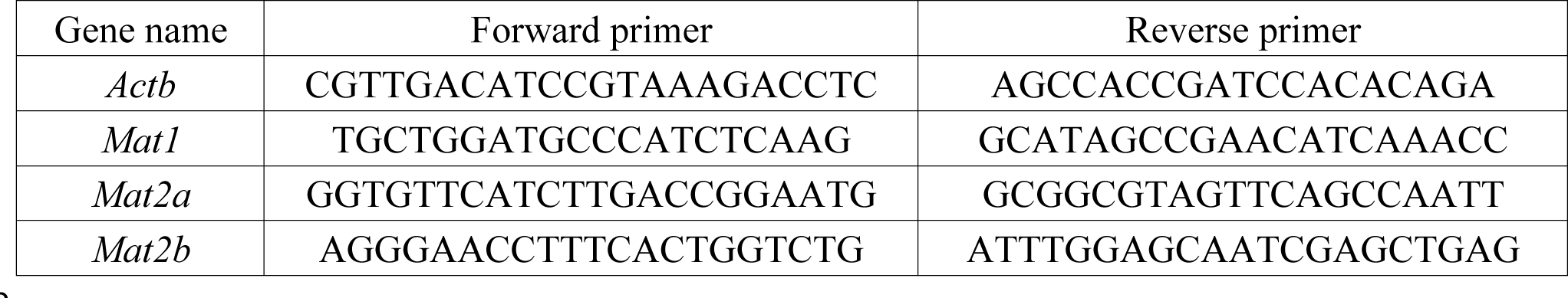
Sequences of the qPCR primers.

### Cell death assessment by flow cytometry

Propidium iodide (PI) and annexin V staining were used for assessment of cell death. Hepa1 or primary hepatocytes were stained by APC-Annexin V according to the manufacturer’s protocols. PI was added (1 µg/mL) before flow cytometry. Hepa1 or primary hepatocytes were sorted with a FACS Verse (BD), and analyzed by FlowJo software (Tree Star, Ashland, OR, USA). Cells that were positive for either or both of annexin V and PI were assessed as dead cells. Conversely, cells that were negative of both annexin V and PI were assessed as alive cells.

### Measurement of endogenous SAM by ELISA kit

S-Adenosylmethionine (SAM) ELISA Kit® (STA-672) was purchased from CELL BIOLABS (San Diego, CA, USA). A Hepa1 cell lysate was prepared according to the attached protocol, and absorbance was measured using a microplate reader iMark^TM^ Microplate Reader (Bio-Rad). Based on the measurement results, a calibration curve was created using Microplate Manager® 6 (Bio-Rad) to calculate the intracellular SAM concentration.

### Statistics

For all experiments, differences of data sets were considered statistically significant when *P*-values were lower than 0.05. Statistical comparisons were performed using the two-sided *t*-test in comparison between the two groups. For the *t*-test, student’s *t*-test was used when the standard deviation (SD) of the groups was not significantly different by *f*-test. Welch’s *t*-test was used when the SD of the groups was significantly different by *f*-test.

### Manuscript preparation

All authors had access to the study data and had reviewed and approved the final manuscript

## List of Abbreviations

IFALD: intestinal failure associated liver disease
MAT: methionine adenosyltransferase
SAM: *S*-adenosylmethionine
ROS: reactive oxygen species

## Author contributions

KT, HN, HS, AM and KI conceived the project, KT carried out the main experiments with support from HN, HS and AM. MW provided suggestions and insights into related diseases. KT, HN, SH, AM and KI interpreted the experimental results, KT wrote the initial manuscript and KI edited. All authors read the manuscript and made comments.

## Data Transparency

Materials related to this manuscript are available upon request.

## Synopsis

Ferroptosis in response to cysteine restriction in hepatoma cells was found dependent on methionine in the media and the syntheses of S-adenosylmethionine and polyamine. Therefore, the status of methionine metabolism may either aggravate or impede ferroptosis under specific conditions.

## References

1. Duggan CP, Jaksic T. Pediatric Intestinal Failure. N Engl J Med. 2017;377(7):666–75.

2. Khalaf RT, Sokol RJ. New Insights Into Intestinal Failure-Associated Liver Disease in Children. Hepatology. 2020;71(4):1486–98.

3. Norsa L, Nicastro E, Di Giorgio A, Lacaille F, D’Antiga L. Prevention and Treatment of Intestinal Failure-Associated Liver Disease in Children. Nutrients. 2018;10(6).

4. Moss RL, Haynes AL, Pastuszyn A, Glew RH. Methionine infusion reproduces liver injury of parenteral nutrition cholestasis. Pediatr Res. 1999;45(5 Pt 1):664–8.

5. Chen J, Li X, Ge C, Min J, Wang F. The multifaceted role of ferroptosis in liver disease. Cell Death Differ. 2022;29(3):467–80.

6. Zhou L, Chen Z, Liu C. Identification and verification of the role of crucial genes through which methionine restriction inhibits the progression of colon cancer cells. Oncol Lett. 2022;24(2):274.

7. Wanders D, Hobson K, Ji X. Methionine Restriction and Cancer Biology. Nutrients. 2020;12(3).

8. Strekalova E, Malin D, Rajanala H, Cryns VL. Preclinical Breast Cancer Models to Investigate Metabolic Priming by Methionine Restriction. Methods Mol Biol. 2019;1866:61–73.

9. Sedillo JC, Cryns VL. Targeting the methionine addiction of cancer. Am J Cancer Res. 2022;12(5):2249–76.

10. Xu Q, Li Y, Gao X, Kang K, Williams JG, Tong L, Liu J, Ji M, Deterding LJ, Tong X, Locasale JW, Li L, Shats I, Li X. HNF4α regulates sulfur amino acid metabolism and confers sensitivity to methionine restriction in liver cancer. Nat Commun. 2020;11(1):3978.

11. Lees EK, Krol E, Shearer K, Mody N, Gettys TW, Delibegovic M. Effects of hepatic protein tyrosine phosphatase 1B and methionine restriction on hepatic and whole-body glucose and lipid metabolism in mice. Metabolism. 2015;64(2):305–14.

12. Obata F, Miura M. Enhancing S-adenosyl-methionine catabolism extends Drosophila lifespan. Nat Commun. 2015;6:8332.

13. Boon R, Kumar M, Tricot T, Elia I, Ordovas L, Jacobs F, One J, De Smedt J, Eelen G, Bird M, Roelandt P, Doglioni G, Vriens K, Rossi M, Vazquez MA, Vanwelden T, Chesnais F, El Taghdouini A, Najimi M, Sokal E, Cassiman D, Snoeys J, Monshouwer M, Hu WS, Lange C, Carmeliet P, Fendt SM, Verfaillie CM. Amino acid levels determine metabolism and CYP450 function of hepatocytes and hepatoma cell lines. Nat Commun. 2020;11(1):1393.

14. Fabris G, Dumortier O, Pisani DF, Gautier N, Van Obberghen E. Amino acid-induced regulation of hepatocyte growth: possible role of Drosha. Cell Death Dis. 2019;10(8):566.

15. Kotb M, Mudd SH, Mato JM, Geller AM, Kredich NM, Chou JY, Cantoni GL. Consensus nomenclature for the mammalian methionine adenosyltransferase genes and gene products. Trends Genet. 1997;13(2):51–2.

16. Murray B, Antonyuk SV, Marina A, Van Liempd SM, Lu SC, Mato JM, Hasnain SS, Rojas AL. Structure and function study of the complex that synthesizes S-adenosylmethionine. IUCrJ. 2014;1(Pt 4):240–9.

17. Shafqat N, Muniz JR, Pilka ES, Papagrigoriou E, von Delft F, Oppermann U, Yue WW. Insight into S-adenosylmethionine biosynthesis from the crystal structures of the human methionine adenosyltransferase catalytic and regulatory subunits. Biochem J. 2013;452(1):27–36.

18. Bailey J, Douglas H, Masino L, de Carvalho LPS, Argyrou A. Human Mat2A Uses an Ordered Kinetic Mechanism and Is Stabilized but Not Regulated by Mat2B. Biochemistry. 2021;60(47):3621–32.

19. Wu C, Jin X, Tsueng G, Afrasiabi C, Su AI. BioGPS: building your own mash-up of gene annotations and expression profiles. Nucleic Acids Res. 2016;44(D1):D313–6.

20. Calvisi DF, Simile MM, Ladu S, Pellegrino R, De Murtas V, Pinna F, Tomasi ML, Frau M, Virdis P, De Miglio MR, Muroni MR, Pascale RM, Feo F. Altered methionine metabolism and global DNA methylation in liver cancer: relationship with genomic instability and prognosis. Int J Cancer. 2007;121(11):2410–20.

21. Mentch SJ, Locasale JW. One-carbon metabolism and epigenetics: understanding the specificity. Ann N Y Acad Sci. 2016;1363(1):91–8.

22. Reytor E, Pérez-Miguelsanz J, Alvarez L, Pérez-Sala D, Pajares MA. Conformational signals in the C-terminal domain of methionine adenosyltransferase I/III determine its nucleocytoplasmic distribution. Faseb j. 2009;23(10):3347–60.

23. Katoh Y, Ikura T, Hoshikawa Y, Tashiro S, Ito T, Ohta M, Kera Y, Noda T, Igarashi K. Methionine adenosyltransferase II serves as a transcriptional corepressor of Maf oncoprotein. Mol Cell. 2011;41(5):554–66.

24. Kera Y, Katoh Y, Ohta M, Matsumoto M, Takano-Yamamoto T, Igarashi K. Methionine adenosyltransferase II-dependent histone H3K9 methylation at the COX-2 gene locus. J Biol Chem. 2013;288(19):13592–601.

25. Biggar KK, Li SS. Non-histone protein methylation as a regulator of cellular signalling and function. Nat Rev Mol Cell Biol. 2015;16(1):5–17.

26. Alam M, Shima H, Matsuo Y, Long NC, Matsumoto M, Ishii Y, Sato N, Sugiyama T, Nobuta R, Hashimoto S, Liu L, Kaneko MK, Kato Y, Inada T, Igarashi K. mTORC1-independent translation control in mammalian cells by methionine adenosyltransferase 2A and S-adenosylmethionine. J Biol Chem. 2022;298(7):102084.

27. Cantoni GL. Biological methylation: selected aspects. Annu Rev Biochem. 1975;44:435–51.

28. Vance DE. Phospholipid methylation in mammals: from biochemistry to physiological function. Biochim Biophys Acta. 2014;1838(6):1477–87.

29. Finkelstein JD. Methionine metabolism in mammals. J Nutr Biochem. 1990;1(5):228–37.

30. Parkhitko AA, Jouandin P, Mohr SE, Perrimon N. Methionine metabolism and methyltransferases in the regulation of aging and lifespan extension across species. Aging Cell. 2019;18(6):e13034.

31. Asai Y, Yamada T, Tsukita S, Takahashi K, Maekawa M, Honma M, Ikeda M, Murakami K, Munakata Y, Shirai Y, Kodama S, Sugisawa T, Chiba Y, Kondo Y, Kaneko K, Uno K, Sawada S, Imai J, Nakamura Y, Yamaguchi H, Tanaka K, Sasano H, Mano N, Ueno Y, Shimosegawa T, Katagiri H. Activation of the Hypoxia Inducible Factor 1α Subunit Pathway in Steatotic Liver Contributes to Formation of Cholesterol Gallstones. Gastroenterology. 2017;152(6):1521–35.e8.

32. Homma T, Kobayashi S, Fujii J. Methionine Deprivation Reveals the Pivotal Roles of Cell Cycle Progression in Ferroptosis That Is Induced by Cysteine Starvation. Cells. 2022;11(10).

33. Li Y, Wang X, Huang Z, Zhou Y, Xia J, Hu W, Wang X, Du J, Tong X, Wang Y. CISD3 inhibition drives cystine-deprivation induced ferroptosis. Cell Death Dis. 2021;12(9):839.

34. Lee J, You JH, Shin D, Roh JL. Inhibition of Glutaredoxin 5 predisposes Cisplatin-resistant Head and Neck Cancer Cells to Ferroptosis. Theranostics. 2020;10(17):7775–86.

35. Stockwell BR. Ferroptosis turns 10: Emerging mechanisms, physiological functions, and therapeutic applications. Cell. 2022;185(14):2401–21.

36. Pope LE, Dixon SJ. Regulation of ferroptosis by lipid metabolism. Trends Cell Biol. 2023;33(12):1077–87.

37. Shima H, Matsumoto M, Ishigami Y, Ebina M, Muto A, Sato Y, Kumagai S, Ochiai K, Suzuki T, Igarashi K. S-Adenosylmethionine Synthesis Is Regulated by Selective N(6)-Adenosine Methylation and mRNA Degradation Involving METTL16 and YTHDC1. Cell Rep. 2017;21(12):3354–63.

38. Pendleton KE, Chen B, Liu K, Hunter OV, Xie Y, Tu BP, Conrad NK. The U6 snRNA m(6)A Methyltransferase METTL16 Regulates SAM Synthetase Intron Retention. Cell. 2017;169(5):824–35.e14.

39. Dai Z, Mentch SJ, Gao X, Nichenametla SN, Locasale JW. Methionine metabolism influences genomic architecture and gene expression through H3K4me3 peak width. Nat Commun. 2018;9(1):1955.

40. Ou Y, Wang SJ, Li D, Chu B, Gu W. Activation of SAT1 engages polyamine metabolism with p53-mediated ferroptotic responses. Proc Natl Acad Sci U S A. 2016;113(44):E6806–E12.

41. Lee BC, Kaya A, Ma S, Kim G, Gerashchenko MV, Yim SH, Hu Z, Harshman LG, Gladyshev VN. Methionine restriction extends lifespan of Drosophila melanogaster under conditions of low amino-acid status. Nat Commun. 2014;5:3592.

42. Yoshida S, Yamahara K, Kume S, Koya D, Yasuda-Yamahara M, Takeda N, Osawa N, Chin-Kanasaki M, Adachi Y, Nagao K, Maegawa H, Araki SI. Role of dietary amino acid balance in diet restriction-mediated lifespan extension, renoprotection, and muscle weakness in aged mice. Aging Cell. 2018;17(4):e12796.

43. Bárcena C, Quirós PM, Durand S, Mayoral P, Rodríguez F, Caravia XM, Mariño G, Garabaya C, Fernández-García MT, Kroemer G, Freije JMP, López-Otín C. Methionine Restriction Extends Lifespan in Progeroid Mice and Alters Lipid and Bile Acid Metabolism. Cell Rep. 2018;24(9):2392–403.

44. Zou K, Rouskin S, Dervishi K, McCormick MA, Sasikumar A, Deng C, Chen Z, Kaeberlein M, Brem RB, Polymenis M, Kennedy BK, Weissman JS, Zheng J, Ouyang Q, Li H. Life span extension by glucose restriction is abrogated by methionine supplementation: Cross-talk between glucose and methionine and implication of methionine as a key regulator of life span. Sci Adv. 2020;6(32):eaba1306.

45. Parkhitko AA, Wang L, Filine E, Jouandin P, Leshchiner D, Binari R, Asara JM, Rabinowitz JD, Perrimon N. A genetic model of methionine restriction extends Drosophila health- and lifespan. Proc Natl Acad Sci U S A. 2021;118(40).

46. Peng H, Yan Y, He M, Li J, Wang L, Jia W, Yang L, Jiang J, Chen Y, Li F, Yuan X, Pang L. SLC43A2 and NFκB signaling pathway regulate methionine/cystine restriction-induced ferroptosis in esophageal squamous cell carcinoma via a feedback loop. Cell Death Dis. 2023;14(6):347.

47. Upadhyayula PS, Higgins DM, Mela A, Banu M, Dovas A, Zandkarimi F, Patel P, Mahajan A, Humala N, Nguyen TTT, Chaudhary KR, Liao L, Argenziano M, Sudhakar T, Sperring CP, Shapiro BL, Ahmed ER, Kinslow C, Ye LF, Siegelin MD, Cheng S, Soni R, Bruce JN, Stockwell BR, Canoll P. Dietary restriction of cysteine and methionine sensitizes gliomas to ferroptosis and induces alterations in energetic metabolism. Nat Commun. 2023;14(1):1187.

48. Hayashima K, Kimura I, Katoh H. Role of ferritinophagy in cystine deprivation-induced cell death in glioblastoma cells. Biochem Biophys Res Commun. 2021;539:56–63.

49. Kan X, Yin Y, Song C, Tan L, Qiu X, Liao Y, Liu W, Meng S, Sun Y, Ding C. Newcastle-disease-virus-induced ferroptosis through nutrient deprivation and ferritinophagy in tumor cells. iScience. 2021;24(8):102837.

50. Nishizawa H, Matsumoto M, Shindo T, Saigusa D, Kato H, Suzuki K, Sato M, Ishii Y, Shimokawa H, Igarashi K. Ferroptosis is controlled by the coordinated transcriptional regulation of glutathione and labile iron metabolism by the transcription factor BACH1. J Biol Chem. 2020;295(1):69–82.

51. Belaidi AA, Masaldan S, Southon A, Kalinowski P, Acevedo K, Appukuttan AT, Portbury S, Lei P, Agarwal P, Leurgans SE, Schneider J, Conrad M, Bush AI, Ayton S. Apolipoprotein E potently inhibits ferroptosis by blocking ferritinophagy. Mol Psychiatry. 2022.

52. Conlon M, Poltorack CD, Forcina GC, Armenta DA, Mallais M, Perez MA, Wells A, Kahanu A, Magtanong L, Watts JL, Pratt DA, Dixon SJ. A compendium of kinetic modulatory profiles identifies ferroptosis regulators. Nat Chem Biol. 2021;17(6):665–74.

53. Funakoshi-Tago M, Sumi K, Kasahara T, Tago K. Critical roles of Myc-ODC axis in the cellular transformation induced by myeloproliferative neoplasm-associated JAK2 V617F mutant. PLoS One. 2013;8(1):e52844.

54. Koomoa DL, Geerts D, Lange I, Koster J, Pegg AE, Feith DJ, Bachmann AS. DFMO/eflornithine inhibits migration and invasion downstream of MYCN and involves p27Kip1 activity in neuroblastoma. Int J Oncol. 2013;42(4):1219–28.

55. Rimpi S, Nilsson JA. Metabolic enzymes regulated by the Myc oncogene are possible targets for chemotherapy or chemoprevention. Biochem Soc Trans. 2007;35(Pt 2):305–10.

56. Casero RA, Jr., Murray Stewart T, Pegg AE. Polyamine metabolism and cancer: treatments, challenges and opportunities. Nat Rev Cancer. 2018;18(11):681–95.

57. Yang WS, Stockwell BR. Ferroptosis: Death by Lipid Peroxidation. Trends Cell Biol. 2016;26(3):165–76.

58. Tsurusaki S, Tsuchiya Y, Koumura T, Nakasone M, Sakamoto T, Matsuoka M, Imai H, Yuet-Yin Kok C, Okochi H, Nakano H, Miyajima A, Tanaka M. Hepatic ferroptosis plays an important role as the trigger for initiating inflammation in nonalcoholic steatohepatitis. Cell Death Dis. 2019;10(6):449.

59. Yamada N, Karasawa T, Kimura H, Watanabe S, Komada T, Kamata R, Sampilvanjil A, Ito J, Nakagawa K, Kuwata H, Hara S, Mizuta K, Sakuma Y, Sata N, Takahashi M. Ferroptosis driven by radical oxidation of n-6 polyunsaturated fatty acids mediates acetaminophen-induced acute liver failure. Cell Death Dis. 2020;11(2):144.

60. Zhou X, Fu Y, Liu W, Mu Y, Zhang H, Chen J, Liu P. Ferroptosis in Chronic Liver Diseases: Opportunities and Challenges. Front Mol Biosci. 2022;9:928321.

61. Tutusaus A, de Gregorio E, Cucarull B, Cristóbal H, Aresté C, Graupera I, Coll M, Colell A, Gausdal G, Lorens JB, García de Frutos P, Morales A, Marí M. A Functional Role of GAS6/TAM in Nonalcoholic Steatohepatitis Progression Implicates AXL as Therapeutic Target. Cell Mol Gastroenterol Hepatol. 2020;9(3):349–68.

62. Bathish B, Robertson H, Dillon JF, Dinkova-Kostova AT, Hayes JD. Nonalcoholic steatohepatitis and mechanisms by which it is ameliorated by activation of the CNC-bZIP transcription factor Nrf2. Free Radic Biol Med. 2022;188:221–61.

63. Jiang L, Wang N, Cheng S, Liu Y, Chen S, Wang Y, Cai W. RNA-sequencing identifies novel transcriptomic signatures in intestinal failure-associated liver disease. J Pediatr Surg. 2022;57(9):158-

